# When Rivals Are Absent: Male Aggression Towards Females in Bluefin Killifish

**DOI:** 10.1101/2024.10.11.617928

**Authors:** Ratna Karatgi, Rebecca C. Fuller

## Abstract

The process of obtaining mates, mating, and (potentially) caring for offspring is costly. While there are inherent costs to reproduction, behavioral interactions among individuals are often the primary drivers of reproductive costs. Males frequently compete for territories and females; females may compete for food or males; males often harass females. Here, we sought to determine whether reproductive costs were primarily due to male/male competition, female/female competition, or male/female interactions in the bluefin killifish. In this species, males guard small spawning territories where females visit them daily to spawn. To manipulate the potential for male and female competition and male/female interactions, we altered the sex ratio and density of each sex across four treatments (1 male: 1 female, 1 male : 3 females, 3 males: 1 female, 3 males: 3 females). Female mortality was higher than male mortality. Surprisingly, female mortality and male aggressive behaviors towards females (i.e., chases) were highest in treatments with a single male. Male-male aggression was present, but males often resolved these disputes via signaling by flaring their fins. There was little evidence for overt aggression among females. When males lack rivals, they turn their territorial defense towards females. These costs help explain why, in nature, females promptly leave male territories following spawning and join loose shoals with conspecific females and minnows.

## Introduction

Reproduction is costly in nearly all taxa. Females produce eggs, and males produce sperm, both of which involve substantial investment of resources. Yet, many of the costs of reproduction stem from behavioral interactions, including aggression (Clutton- Brock and Parker, 1995; Chapman *et al*., 2003; Wedell *et al*., 2006; Pizzari and Bonduriansky, 2010). Males often compete with one another over access to mating territories or females themselves (Emlen and Oring, 1977). In some systems, females compete for access to males (Clutton-Brock, 2007; Rosvall, 2011; Stockley and Campbell, 2013). Sexual conflict between males and females often results in costs for one or both sexes (Arnqvist and Rowe, 1995; Pizzari and Bonduriansky, 2010; Makowicz and Schlupp, 2013). Teasing apart the effects of agonistic behaviors during reproduction can be complex, but is necessary to understand the selective forces acting on males and females and the resulting mating system(Emlen and Oring, 1977; Kvarnemo and Ahnesjö, 1996; Wedell *et al*., 2006).

Manipulations of the operational sex ratio are often informative to understand the nature of sexual selection emanating from male competition, female competition, and male/female interactions (De Jong *et al*., 2012). The operational sex ratio is defined as the ratio of males to females ready to mate at a given time (Clutton-Brock and Parker, 1992; Kvarnemo and Ahnesjö, 1996; Ahnesjö *et al.,* 2001). Male-biased sex ratios (more males ready to mate than females) are thought to promote competition among males over females. Indeed, several studies have found that as the number of reproductively available females decreases, the intensity of male intrasexual competition over mating opportunities increases (Schwagmeyer and Brown, 1983; Colwell and Oring, 1988; Kodric-Brown, 1988; Tejedo, 1988; Kvarnemo *et al.,* 1995; Cureton *et al.,* 2010; Wacker *et al.,* 2013; Chuard *et al.,* 2016). Variations in environmental conditions, such as ambient temperature and spawning substrate quality, which make the operational sex ratio more male-biased, also lead to increased male intrasexual competition (Zamudio *et al.,* 1995; Forsgren *et al.,* 1996; Kvarnemo, 1996; Forsgren *et al.,* 2004). In some species, the density of conspecifics, rather than the operational sex ratio, has a significant effect on male competition and aggression (Jirotkul, 1999) In recent years, competition among females has received more empirical support(Clutton-Brock, 2007, 2009; Rosvall, 2011 and references within). Traditionally, competition among females was thought to be confined to sex-role-reversed species where males are the limiting resource in reproduction, males are choosy among females, and females compete for males (Colwell and Oring, 1988; Berglund, 1994; Ahnesjo, 1995). Yet, new evidence has found pronounced competition among females in systems with ‘traditional’ sex roles where females are the limiting resource in reproduction (Rosvall, 2011; Ranade *et al*., 2024).

Finally, reproductive costs often emerge due to behavioral interactions between males and females. Differences between males and females in the optimal mating rates result in the sexes having different optima for various traits, leading to sexual conflict (Trivers, 1972; Rice, 1996; Chapman *et al*., 2003). Male mate monopolization not only leads to intense competition between males but can also result in female harm by males. In species with internal fertilization, this can manifest as physical harm to female reproductive tracts, as seen in birds and many insects (Waage, 1979; Arnqvist and Rowe, 1995; Crudgington and Siva-Jothy, 2000), or through proteins present in seminal fluid that can reduce female lifespan, as seen in Drosophila (Chapman *et al.,* 1995; Kuijper *et al.,* 2006; Chapman, 2018). An indirect effect of male harassment is that females spend more time avoiding male-dominated regions, which can lead to a reduction in time spent foraging (Arnqvist, 1989; Krupa and Sih, 1993; Rowe, 1994; Schlupp *et al.,* 2001; Pilastro *et al.,* 2003; Plath *et al.,* 2003) or an increase in exposure to predators (Arnqvist, 1989; Rowe, 1994; Darden and Croft, 2008). As a result, females often pay a high cost due to male coercion, which can be seen in the form of a reduction in fecundity (McLain and Pratt, 1999; Rossi *et al.,* 2010), body condition (Watson et al., 1998; Makowicz and Schlupp, 2013), and, in extreme situations, mortality (Mcentee *et al*., 2023). Even in lek mating species where males cannot control access to females, male harassment of females is seen and can have dire consequences for females (Reale *et al.,* 1996). Male mate harm is typically most intense during the periods of the breeding season when the sex ratio is highly male-biased (Reale *et al.,* 1996; Holand *et al.,* 2006). Although male mate harassment has been documented in a broad range of taxa, much of this research is on internal fertilizers. Male mate harm in external fertilizers, including many fish and amphibians, has received less attention. In these species, male harassment of females is more likely to be through direct physical aggression rather than through mechanisms during copulation or post-copulatory seminal fluids.

In this study, we manipulate the operational sex ratio to examine the roles of male-male, female-female, and male-female interactions on aggression and mortality in the bluefin killifish, *Lucania goodei*, a freshwater fundulid fish. In this species, males defend small territories by flaring their fins, chasing, and attacking neighboring males (Foster, 1967; Arndt, 1971). Males quickly establish a dominance hierarchy that remains stable for a relatively long duration (McGhee and Travis, 2010). When a female enters a male’s territory, the male either courts the female or chases her away (Fuller, 2001). During courtship, males swim in circles around the female, flash their fins, and flick their heads in the vicinity of the female (Foster, 1967; Arndt, 1971; Breder and Rosen, 1996). If the female accepts the male, they proceed to pair spawn on aquatic vegetation (Fuller, 2001); however, in laboratory settings, the most active males spawned with females, overriding female choice (McGhee *et al.,* 2007). Following spawning, the female leaves the territory and joins loose shoals containing conspecific females, non-territory holding males, and heterospecific fish (i.e., minnows). Although males guard their territories and are often aggressive well after spawning, there is no strong evidence of male parental care towards the clutches spawned in their territories (Fuller and Travis, 2001). In bluefin killifish, the spawning season is quite long, extending from February through September (Fuller, 2001), with some authors reporting fish that breed year-round (Arndt, 1971). Females are highly iteroparous (Arndt, 1971; Breder and Rosen, 1996). Females become gravid multiple times each breeding season and spawn daily for about two weeks when gravid.

Our specific goals were to determine whether the number of females, number of males, and sex ratio influence sex-specific aggression and mortality. We ask the following questions: (a) Does male intrasexual aggression vary between male-biased and even sex ratio treatments? (b) Do females show intrasexual aggression? If so, does it differ between female-biased and even sex ratio treatments? (c) Does male aggression towards females vary as a function of sex ratio and density treatments? (d) Does male and female survivorship vary as a function of sex ratio and density?

## Methods

This experiment aimed to assay the levels of aggression in male and female bluefin killifish (both within and between the sexes) and to determine the extent to which aggression led to detrimental effects on survival. We manipulated the sex ratio and density of bluefin killifish in 110-liter tanks and observed behavior and survival.

Male and female bluefin killifish (*Lucania goodei*) were caught using seine nets from the Wakulla River, Florida. Fish were transported back to the University of Illinois, where they were housed in a greenhouse in 400-liter stock tanks with large sponge filters that aerated the water and removed nitrogenous wastes. Fish were fed brine shrimp daily.

We manipulated sex ratios and density to create treatments where competition and aggression were likely to differ between males and females. We created a treatment with a male-biased sex ratio (3 males: 1 female) and a female-biased sex ratio (1 male: 3 females). We also had two controls with an even sex ratio to control for the density of each sex (1 male : 1 female and 3 males : 3 females). Six replicates were performed across the four treatments (24 tanks in total). The experiment was carried out between February 2020 – July 2020. The experimental tanks were initially set up in late February 2020. Due to the COVID-19 pandemic, they remained in those conditions for six months. During this time, tanks were censused, fish mortality was noted, and dead fish were replaced. We measured survival as the proportion of individuals alive in each treatment, which was the total number of individuals of each sex alive divided by the total number placed in the tank.

We conducted behavioral observations from June - to July 2020. Fish were marked using a fluorescent elastomer dye injected underneath their skin, enabling individual identification within a tank (Northwest Marine Technology, Inc.). Each fish in each tank was observed once for a duration of five minutes. During these observations, we recorded the number of chases and attacks exhibited by both males and females. The number of fin flares and courtship bouts was also noted for males. For chases and attacks, we recorded the sex of the recipient of aggression by the focal individual.

Behavioral observations were recorded using voice recordings, and the resultant audio files were transcribed using BORIS (Friard and Gamba, 2016).

### Statistical Analysis

*Survival.*- We measured the survival of both males and females for each tank. Animals died over the course of this experiment, and we replaced them. The survival of each sex was the total number of animals that lived divided by the total number of animals placed in that tank. We first asked whether survival differed between males and females. We used a mixed model where we examined the effect of sex on survival, treating individual tanks as a random effect. We found a significant effect of sex and therefore analyzed the survival of each sex separately. We performed a generalized linear model for male and female survival, examining the effects of female density (1 or 3), male density (1 or 3), and their interaction. We specified a binomial distribution for all three models. We checked for over-dispersion for each model but found none. All analyses of variance used type 3 comparisons and were conducted using the ‘car’ package in R. The options for all analyses were set to *options(contrasts = c(“contr.sum”, “contr.poly”))*.

*Aggression and Signaling. -* This experiment aimed to examine aggression within each of the sexes. Female aggression was rare and observed only in one instance. Therefore, we analyzed only male aggression and signaling behaviors. We compared two measures of male-male aggression (attacks and chases) between the two treatments where the number of males was 3 (3m1f and 3m3f). For both behaviors, we calculated the mean for each tank. Attacks were log-transformed to meet the assumptions of analysis of variance (normal residuals, no heteroscedasticity). We performed a linear model where we compared the levels of aggression between the two treatments (3m1f and 3m3f).

Male signaling was measured as the number of fin flares given by a male. Males flared their fins towards both males and females. We measured the mean number of fin flares performed for each tank. We log-transformed the mean number of fin flares given to meet the assumptions of the analysis of variance. Post hoc comparisons were made using Tukey’s HSD from the emmeans package.

Male aggression towards females was in the form of chases and attacks. We analyzed the effects of the different treatments on mean and mean per capita aggression experienced by females. Intersexual aggression experienced per capita females was calculated as the mean number of aggressive behaviors divided by the number of females in each treatment. Using a generalized linear model, we analyzed the effects of female density (1 or 3), male density (1 or 3), and their interaction on mean male chases and attacks towards females. Since the analysis was on non-integer response variables that deviated significantly from normality and were highly over-dispersed, we specified a gamma distribution for all four models of male aggression towards females. Post hoc comparisons were made using Tukey’s HSD from the emmeans package. All analyses were performed in R (version 4.4.0). Raw data and R codes will be made available at Dryad (link to be inserted upon acceptance).

## Results

*Survival. -* Male survival was significantly higher than female survival (χ^2^= 4.806, p = 0.028, figure 1). Female survival was greatly reduced in treatments where there was one male in comparison to three males (χ^2^ = 6.315, p = 0.012, table 1). Female survival tended to be higher in treatments with three females in the tank compared to one, but the result was not statistically significant (χ^2^ = 2.317, p = 0.128, table 1). The interaction between the number of males and the number of females did not account for a significant amount of variation in female survival (χ^2^= 0.080, p = 0.778). There were no effects of treatment on male survival (p>0.13 in all tests, table 1).

**Figure 1.**
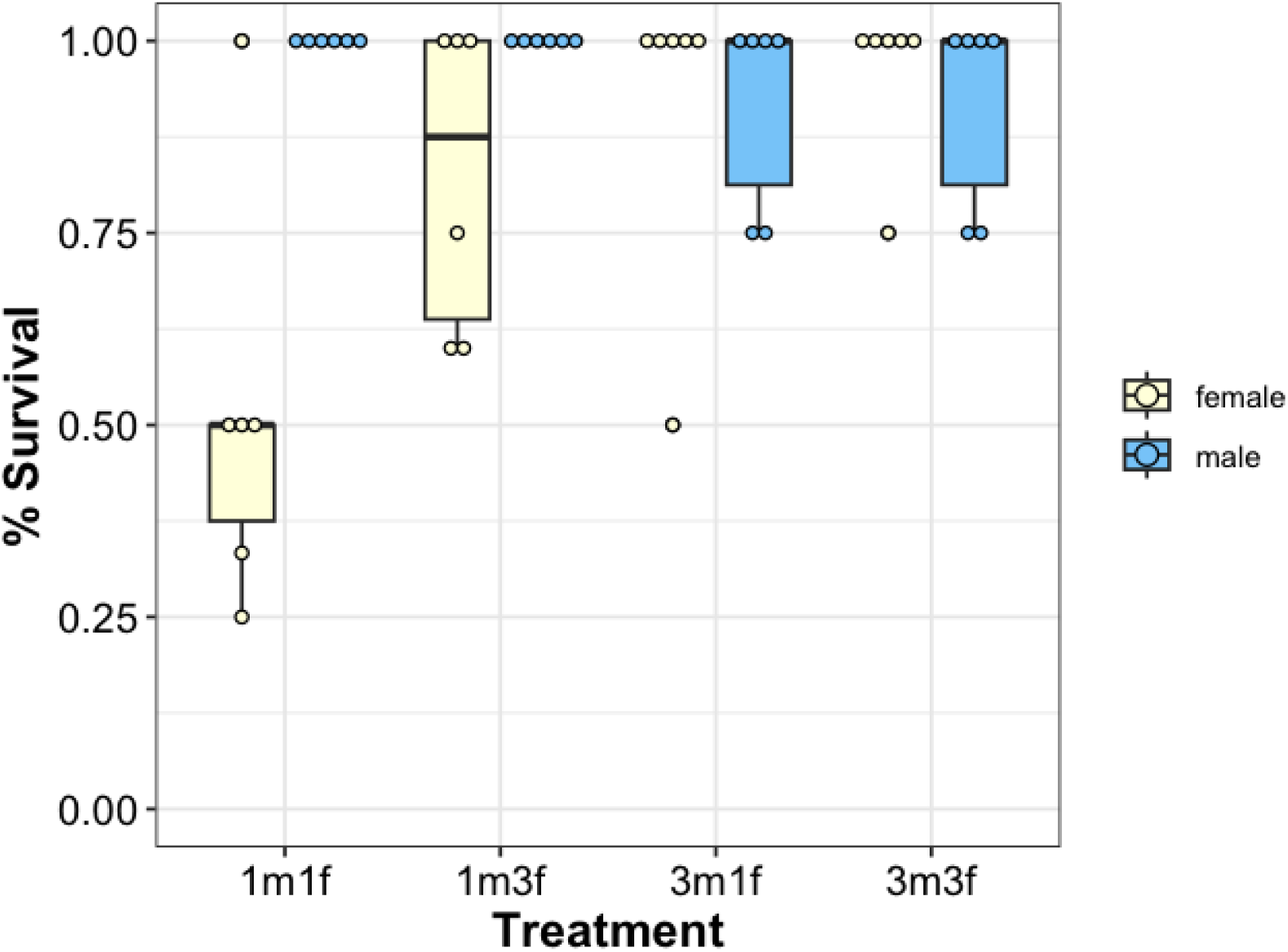
Box plots and raw data for survival for males and females across the four treatments. The number of each sex is given on the X-axis. Survival was measured as the number of animals alive at the end of the experiment divided by the total number of animals put in the tank, including replacement animals.

**Table 1.** Type 3 analysis of deviance on the proportion of (A) females and (B) males alive across the four treatments.

| Terms | $\chi^2$ | df | P |
| --- | --- | --- | --- |
| *A. Female survival* |  |  |  |
| Number of males | 6.31 | 1 | 0.012 |
| Number of females | 2.32 | 1 | 0.128 |
| Number of males x Number of females | 0.08 | 1 | 0.778 |
| *B. Male survival* |  |  |  |
| Number of males | 2.20 | 1 | 0.138 |
| Number of females | 0.00 | 1 | 1.000 |
| Number of males x Number of females | 0.00 | 1 | 1.000 |

*Aggression. -* Females were rarely aggressive, and intrasexual female aggression was observed only in one instance across all the behavioral observations. Although males were aggressive, male intrasexual aggression did not vary as a function of the sex ratio between the 3m1f and 3m3f treatments (Attacks: *F*_1,10_ = 0.693, *p* = 0.424, Chases: *F_1,10_* = 0.712, *p* = 0.422, figure 2).

**Figure 2.**
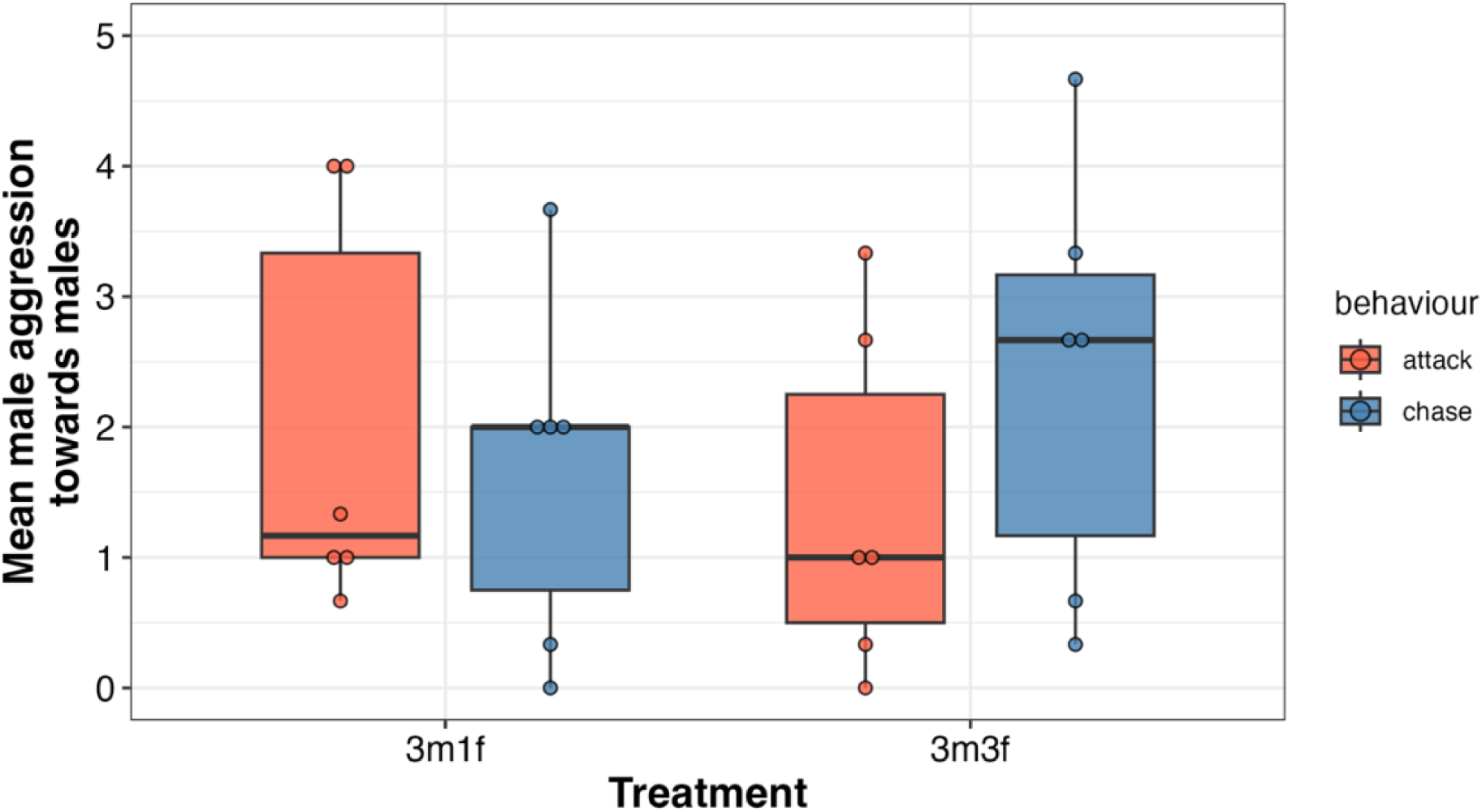
Box plots for the mean number of attacks and chases given by males. Each dot is the mean for one tank. The number of each sex is given on the X-axis.

Male fin flares varied significantly due to the number of males (*F_1,20_*= 11.5075, *p* = 0.0029, figure 3, table 2) but not the number of females or their interaction. Males signaled significantly more in the three male treatments (3m1f, 3m3f) compared to the one male treatments (1m1f, 1m3f).

**Figure 3.**
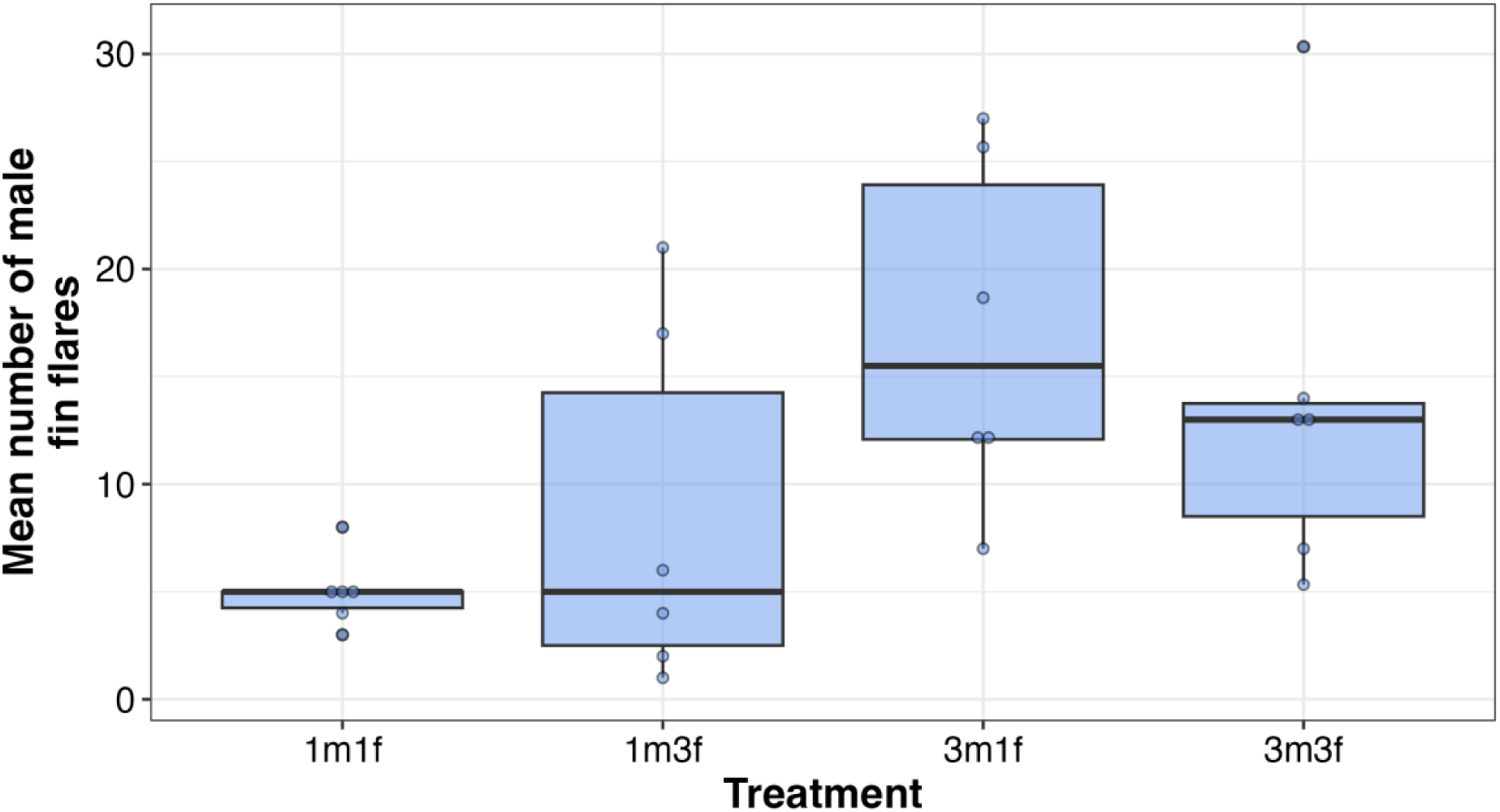
Box plots and mean numbers of fin flares by males across the four treatments. The number of each sex is given on the X-axis.

**Table 2.** Type 3 analysis of variance on the mean number of fin flares performed by males across all four treatments.

| Terms | *F* | *df* (num, denom) | *P* |
| --- | --- | --- | --- |
| Intercept | 317.96 | 1,20 | 0.000 |
| Number of males | 11.51 | 1,20 | 0.003 |
| Number of females | 0.055 | 1,20 | 0.817 |
| Number of males x Number of females | 0.54 | 1,20 | 0.470 |

Male aggression towards females in the form of chases was significantly influenced by the number of males (χ^2^ =7.4656, *p*=0.0063, figure 4a, table 3a) and the number of females (χ^2^ =8.5523, *p*=0.0035), but not their interaction (χ^2^ =0.9358, *p*=0.3334).

**Figure 4.**
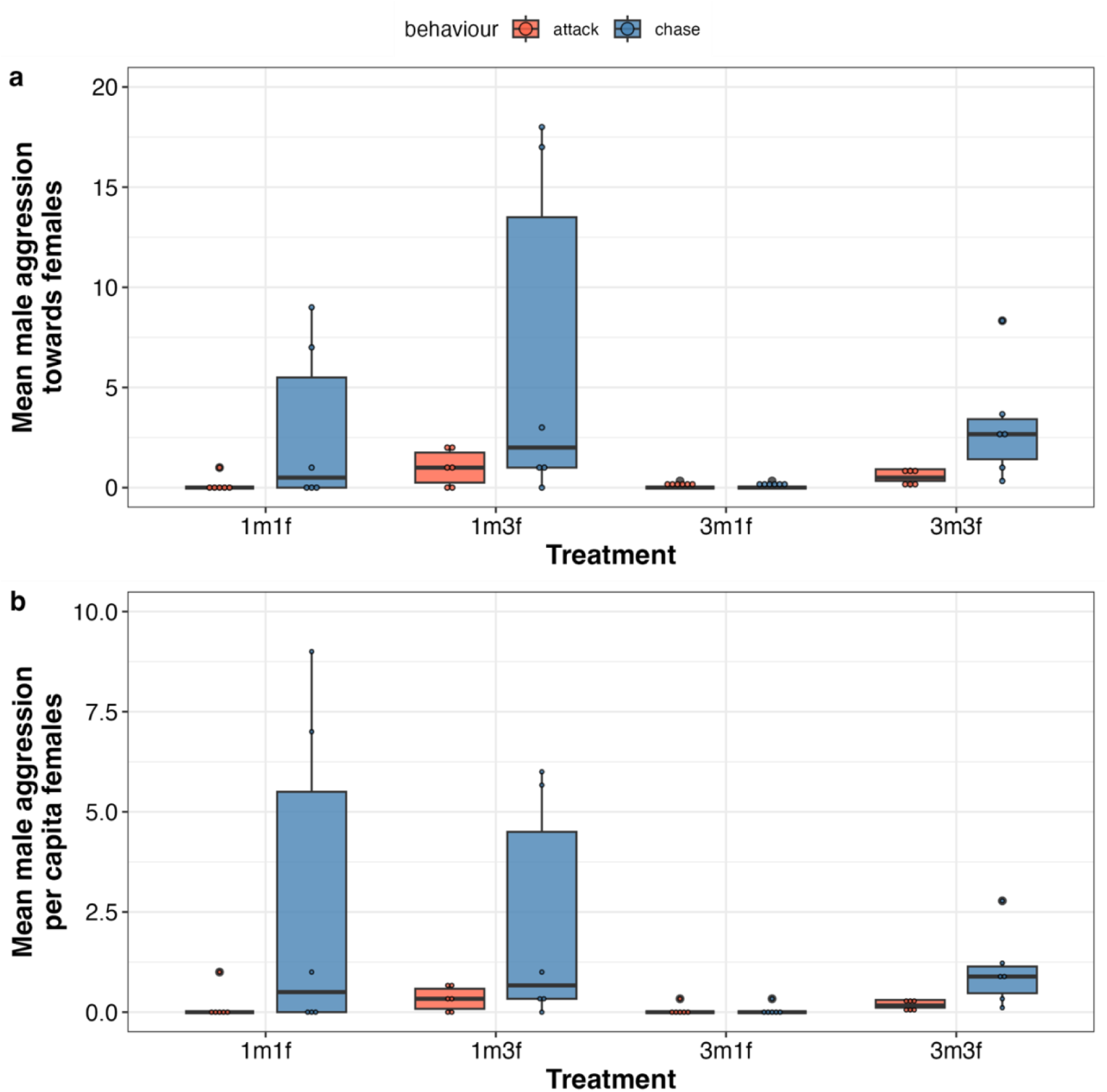
Box plots for the mean number of aggressive behaviors towards females by males across the four treatments. Each dot is an average for each tank. The number of each sex is given on the X-axis. (a) - mean male aggression towards females. (b) – Male aggression per capita female.

**Table 3.** Type 3 analysis of deviance on the mean (A & B) and per capita (C & D) mean aggressive behaviors (chases and attacks) performed by males towards females across all four treatments.

| Terms | $\chi^2$ | df | P |
| --- | --- | --- | --- |
| *A. Mean Chases* |  |  |  |
| Number of males | 7.47 | 1 | 0.006 |
| Number of females | 8.55 | 1 | 0.003 |
| Number of males x Number of females | 0.94 | 1 | 0.333 |
| *B. Mean Attacks* |  |  |  |
| Number of males | 1.82 | 1 | 0.177 |
| Number of females | 12.57 | 1 | 0.0004 |
| Number of males x Number of females | 0.34 | 1 | 0.561 |
| *C. Per capita mean chases* |  |  |  |
| Number of males | 8.37 | 1 | 0.004 |
| Number of females | 0.66 | 1 | 0.416 |
| Number of males x Number of females | 1.94 | 1 | 0.164 |
| *D. Per capita mean attacks* |  |  |  |
| Number of males | 1.41 | 1 | 0.235 |
| Number of females | 1.85 | 1 | 0.174 |
| Number of males x Number of females | 0.009 | 1 | 0.923 |

Attacks towards females were much rarer, and male attacks towards females was significantly influenced by the number of females (χ^2^ =12.5681, *p*=0.00039, table 3b), but not the number of males (χ^2^ =1.8206, *p*=0.177) or the interaction of the two (χ^2^ =0.337, *p*=0.5612). Not surprisingly, males were more aggressive towards females in the treatments with three females (1m3f, 3m3f). In essence, there were more females to be aggressive towards. In some replicates with a single female, we observed females hiding from males. However, contrary to expectations, females experienced greater aggression from males in treatments with one male (1m1f, 1m3f) compared to three males (3m1f). More males did not lead to more aggression towards females.

Male aggression per capita females in the form of chases was significantly influenced by the number of males (χ^2^= 8.3727, *p*= 0.0038, figure 4b, table 3c) but not the number of females (χ^2^= 0.6595, *p*= 0.4167) or their interaction (χ^2^= 1.9374, *p*= 0.1639). Attacks towards females were much rarer, and there were no patterns in male attacking behavior per capita females across the different treatments (number of males -χ^2^ =1.4095, *p*=0.2351, number of females -χ^2^ =1.8462, *p*= 0.1742 and their interaction -χ^2^ =0.0093, *p*=0.9231, table 3d).

## Discussion

In this study, we tested for the roles of density and sex ratio on aggression, signaling, and survival in male and female bluefin killifish. Surprisingly, we found high levels of female mortality when females were housed with a single male. Male aggression towards females was also higher when a single male was housed with multiple females. There was aggression among males when multiple males were housed together, but the number of females had little influence on these levels. Males also signaled via fin flares more when they were with other males, a finding that is in keeping with past work showing that fin flares precede aggression (Arndt, 1971; Fuller, 2001; McGhee *et al.,* 2007). Below, we discuss the implications of these results.

The most surprising finding is that female mortality was high when females were housed with a single male, and this was particularly so when a single female was housed with a single male. In fact, in some tanks, males were particularly aggressive and had to be replaced lest they continue to kill females. In the one male, three female treatments, males often directed their aggression towards specific individual females while ignoring others. These results are surprising because we expected that aggression towards females would be highest in the treatments with three males, particularly three males and one female. In fact, previous studies on other species have shown that the intensity of male harassment toward females tends to decrease as the number of females increases and sex ratio becomes less male-biased (Krupa and Sih, 1993; Vepsäläinen and Savolainent, 1995; Doutrelant *et al.,* 2001; Cureton *et al.,* 2010).

Extensive research has shown that mating is costly for females, but much of this work has focused on insects, birds, and mammals, where internal fertilization inherently imposes greater costs on females (Wing *et al.,* 1983; Crudgington and Siva-Jothy, 2000; Wedell *et al.,* 2006). As a result, we have a deep understanding of the physical and post-copulatory costs females face, including genital tract injuries, the antagonistic effects of seminal fluid proteins, and the consequent evolutionary arms race surrounding reproductive traits (Arnqvist and Rowe, 1995; Chapman *et al*., 1995; Arbuthnott *et al*., 2014; Chapman, 2018). However, these findings do not fully explain how sexual conflict unfolds in species that rely on external fertilization, where the dynamics of mating and reproductive costs may differ significantly.

In this experiment, females were housed continuously with at least one territorial male. An argument can be made that this is artificial because, in nature, females promptly leave the spawning territories to shoal with other females and minnows. We argue that our results shed light on this behavior in the wild; we argue that females leave male territories precisely because further interactions with territorial males are highly costly.

Studies on mosquitofish have found that females experienced reduced male harassment and increased foraging efficiency when they associated with a shoal (Pilastro *et al.,* 2003) and actively chose to be in close proximity to other females in the presence of a harassing male (Dadda *et al.,* 2005). They also preferred to associate with larger females who were more likely to be targeted by a courting male (Agrillo *et al.,* 2006). Non-receptive female Trinidadian guppies chose to associate with receptive over non-receptive females in order to receive lower male agonistic behaviors (Brask *et al*., 2012). In nature, bluefin killifish male aggression towards other males is high and primarily directed towards neighboring males. Aggression towards females is present but much lower because females can leave the male territories (Fuller, 2001), and the results of this study indicate that male aggression can influence female association patterns in the wild.

An outstanding question is why males exhibit such excessive aggression, with several hypotheses still untested. We have frequently observed male-female interactions in the absence of competing males and noted that male aggression towards females spikes after spawning. While several species in the order *Cyprinodontiformes* are known to display male parental care (Breder and Rosen, 1996; Mank *et al.,* 2005), previous studies investigating male parental care, such as increased egg survival, found little evidence of this in bluefin killifish (Fuller and Travis, 2001). Previous work compared egg survival rates in tanks with no fish, males only, and males plus minnows, but it did not specifically examine male behavior toward females in relation to nest guarding.

Importantly, it did not test for egg survival in the presence of males only versus males plus females. We have frequently observed females eating eggs on mops, raising the possibility that male aggression could be a form of nest guarding to protect eggs from female predation.

A second hypothesis is that male aggression towards females represents the effects of spillover of male intrasexual aggressive behavior. In nature, reproductive males spend much of their days defending their territories from neighboring males, and male aggression towards other males is tightly correlated with male mating success. This is also seen in laboratory experiments where male aggression and activity were the most significant predictors of spawning success, overriding the role of female choice (McGhee *et al.,* 2007). In the absence of competing males, males may improperly direct their aggression towards females. In our lab, we have higher female survival in tanks with a single male-female pair when the males could see other males in neighboring tanks. Under these conditions, males tended to direct much of their attention towards their neighbors (Fuller, pers obs).

We observed that male aggression toward other males was lower than aggression directed at females, contrasting with patterns typically seen in the wild (Fuller, 2001). While males did exhibit aggression toward each other, their interactions often involved signaling from a distance rather than direct physical conflict. For example, fin flaring -commonly associated with aggressive behavior in fish (Foster, 1967) - was more frequently used during male-male interactions than male-female aggression. This suggests that males have developed strategies such as initiating aggressive encounters through behaviors as signaling through which they can assess the resource-holding potential of their competitors before engaging in physical conflict, thereby reducing the costs of aggressive encounters (Arndt, 1971; Parker, 1974; van Staaden *et al.,* 2011).

Similarly, females appear to mitigate the likelihood of aggression by distancing themselves from males and forming schools with other females.

In this study, we find that females rarely exhibit aggression, either towards other females or males. Recent research has shown strong evidence of female intrasexual aggression in species without sex-role reversal - whether through competition for mates or interference competition over resources such as food or nest sites (Clutton-Brock, 2009; Rosvall, 2011; Ranade *et al*., 2024). In contrast, our findings suggest that female aggression may be less prominent, particularly in external fertilisers without significant parental care. It is possible that the limitations of our study may have reduced our ability to detect female competition. However, the consistent absence of aggression across all replicate tanks indicates that aggressive behaviors are infrequent among bluefin killifish females in this context. This pattern has also been noted in subsequent experiments unrelated to this study. One explanation for the low levels of female intrasexual aggression could be the prolonged breeding season in bluefin killifish, allowing females ample time with preferred males and making competition for access to males less critical. Additionally, females often shoal with other fish when not in male territories, where aggressive behavior may not be advantageous or necessary.

In summary, our findings indicate that females incur significant reproductive costs due to male territorial behaviors, particularly when no competitor males are present. These costs are highlighted by the heightened aggression females face, which may contribute to reduced female survival in the presence of a single male. This pattern helps to explain observed female movement behaviors in the wild, where females typically avoid prolonged contact with territorial males. By highlighting the reproductive costs associated with mating in an iteroparous, external fertilizer species, our study enhances the understanding of the consequences of sexual conflict in species with external fertilization.

## Author Contributions

Ratna Karatgi: Conceptualization (equal); Investigation (lead); Methodology (equal); Formal analysis (lead); Writing – original draft (lead); Writing - review & editing (lead).

Becky Fuller: Conceptualization (equal); Resources (lead); Methodology (equal); Formal analysis (supporting); Writing - original draft (supporting), Writing – review and editing (supporting).

## Acknowledgements

We thank Evi Malone, Jonny Chan, Kieran Patel, and Jacob Grandison for their help with fish husbandry and experimental setup. The work was approved by the University of Illinois Institutional Animal Care and Use Committee (17184).

## Data Accessibility Statement

All analyses were conducted in R version 4.4.4. The raw data and associated R-code will be uploaded to Dryad upon acceptance.

